# Alpha and theta dynamics successively resolve competition between rival visual stimuli

**DOI:** 10.1101/2025.07.21.665846

**Authors:** Wen Wen, Yichen Wu, Robert M.G. Reinhart, Sheng Li

**Affiliations:** School of Psychological and Cognitive Sciences, Peking University, Beijing, China; PKU-IDG/McGovern Institute for Brain Research, Peking University, Beijing, China; Key Laboratory of Machine Perception (Ministry of Education), Peking University, Beijing, China; Beijing Key Laboratory of Brain-Computer Interface and Mental Health Modulation, Peking University, Beijing, China; Department of Psychological & Brain Sciences, Boston University, Boston, MA 02215, USA

**Keywords:** distractor inhibition, feature-based attention, frontal theta, parietal alpha, binocular rivalry, consciousness

## Abstract

Attention plays a central role in selecting information for conscious perception. Because natural environments are structured, prior knowledge about relevant and irrelevant features can guide this selection. However, prior knowledge about distractors can enhance their neural representation, raising the question of how the brain prevents such enhancement from interfering with behavior. We addressed this question by combining a binocular rivalry paradigm with frequency-tagged stimuli to independently track target and distractor signals during sustained perceptual competition. We identified complementary alpha and theta dynamics that act successively to stable perception. Early parietal alpha activity (8-12 Hz) was associated with the segregation of competing inputs without altering sensory gain. This was followed by frontal theta activity (3-7 Hz), which was linked to the reactive suppression of distractors representations. Importantly, alpha-mediated gating effects were most pronounced under high perceptual uncertainty when sensory evidence were weak, where they facilitated the stabilization of weak target signals and promoted their access to conscious awareness. Together, these findings demonstrate that perceptual competition is resolved through a coordinated sequence of proactive gating and reactive suppression.

## Introduction

Our brain is constantly bombarded with more sensory input than it can fully process, yet we perceive a seamless, unified world rather than a chaotic flux of signals. This stability depends on the brain’s ability to orchestrate access to consciousness, prioritizing goal-relevant inputs while suppressing irrelevant information. Selective attention serves as the primary gatekeeper in this process, resolving the competition for limited processing resources. Importantly, prior experience and learned expectations allow the brain to anticipate upcoming demands and allocate processing resources accordingly (Anderson 2021). Understanding how such knowledge shapes attentional control is therefore fundamental to explaining how the brain constructs a coherent visual reality.

Extensive research has shown that prior knowledge about target features, such as knowing where or what to look for, can significantly enhance attentional performance by facilitating the selection of task-relevant input (Corbetta and Shulman 2002; Gazzaley and Nobre 2012). However, the converse remains less well understood: can attentional selection be improved by knowing what to ignore? This question is particularly important for two key reasons. First, natural environments are structured and governed by regularities. We typically engage with visual scenes informed by prior knowledge derived from working memory (Soto et al. 2008), long-term memory (Hutchinson and Turk-Browne 2012), or learning (Theeuwes et al. 2022). Second, targets rarely appear in isolation. Given the limited computational resources, the brain must arguably do more than just amplify targets; it must actively shield conscious processing from predictable interference. This raises a central question: can the brain utilize distractor knowledge to bar specific features from higher-level processing?

When distractor features are known in advance, the exclusion of irrelevant inputs can occur either proactively before stimulus onset or reactively after interference occurred (Liesefeld, Heinrich R. et al. 2024; Theeuwes 2025). These two forms of inhibition are thought to rely on distinctive neural mechanisms (Chelazzi et al. 2019; Geng 2014; Noonan et al. 2018; van Moorselaar and Slagter 2020). Proactive inhibition acts as a preemptive strategy to suppress distractors and reduce their ability to capture awareness. Studies have shown decreased excitability of corresponding sensory cortex in anticipation of distractors (Ferrante et al. 2023; Snyder and Foxe 2010), as well as stronger suppression of spatial locations where distractors frequently appear (Wang et al. 2019). Reactive inhibition, by contrast, is engaged after distractors are encountered and operates through mechanisms such as disengagement (Liesefeld, Heinrich René et al. 2017; Lin et al. 2024; Moher and Egeth 2012) or signaled suppression (Gaspelin and Luck 2018; Sawaki and Luck 2010). Both forms of inhibition rely on top-down control, mediated by frontal theta activity (3-7Hz), which supports the dynamic regulation of sensory input in accordance with task demands (Cavanagh and Frank 2014; de Vries et al. 2019; Wen et al. 2022).

An unresolved issue is how these inhibitory signals prevent distractors from disrupting visual coherence (Noonan et al. 2018; van Moorselaar and Slagter 2020). One class of accounts proposes that inhibition operates directly by reducing sensory gain for distractor representations. Another class emphasizes indirect control mechanisms, in which attentional priorities are enforced by regulating the flow of information without altering sensory gain. This distinction is closely linked to ongoing debates about the functional role of alpha-band oscillations (8-12 Hz) (Bonnefond and Jensen 2024; Klimesch 2012; Peylo et al. 2021; Yang et al. 2024). Alpha oscillations have long been associated with modulating neuronal excitability of sensory regions and neural responses to sensory input (Foxe and Snyder 2011; Iemi et al. 2022; Richter et al. 2025; Van Diepen et al. 2019). Yet accumulating evidence suggests that alpha oscillations may instead gate information flow in a goal-dependent manner, routing relevant and irrelevant signals differently to determine their access to higher-order processing, without directly suppressing sensory responses (Antonov et al. 2020; Bonnefond and Jensen 2024; Duecker et al. 2025; Gundlach et al. 2020; Morrow et al. 2023; Zhigalov and Jensen 2020). Distinguishing between direct gain modulation and indirect gating therefore remains a central challenge for understanding the neural mechanisms of distractor inhibition.

Here, we addressed this challenge using a binocular rivalry paradigm combined with frequency-tagged steady-state visual evoked responses (SSVERs). Conventional visual search paradigms present a target among multiple distractors within the same visual field, posing several challenges for dissociating their neural processing. The numerical imbalance can complicate direct comparisons between target- and distractor-related neural representations because their aggregate sensory contributions are not matched. Target and distractor signals can also be difficult to separate because of their spatial proximity. Moreover, the competition is relatively brief because it is substantially reduced once the target is identified. Binocular rivalry provides a complementary approach: presenting two incompatible stimuli separately to the two eyes generates strong, sustained competition between spatially overlapping visual inputs. By tagging the two stimuli at different frequencies, we could therefore independently track target-and distractor-related sensory responses throughout ongoing competition. This design allowed us to ask how advance knowledge of either a target or a distractor shapes their subsequent neural processing. Participants viewed two colored gratings flickering at different frequencies, one designated as the target and the other as the distractor (24 Hz for the target and 20 Hz for the distractor; Figure. 1A). Their task was to report the color of the target grating at the end of each trial. On each trial, participants were cued in advance about the orientation of either the upcoming target or distractor grating. In the target-cueing condition (TgtCue), the cue indicated the orientation of the upcoming target, whereas in the distractor-cueing condition (DistCue), it indicated the orientation of the upcoming distractor, allowing participants to identify the target as the grating with the alternative orientation. Thus, both cueing conditions provided task-relevant advance information but differ in its content, allowing us to compare how these two forms of prior knowledge shape attentional control during subsequent visual competition. For each block, participants were informed of the cueing condition ahead (see Methods for discussion of a no-cue condition).

**Figure 1:**
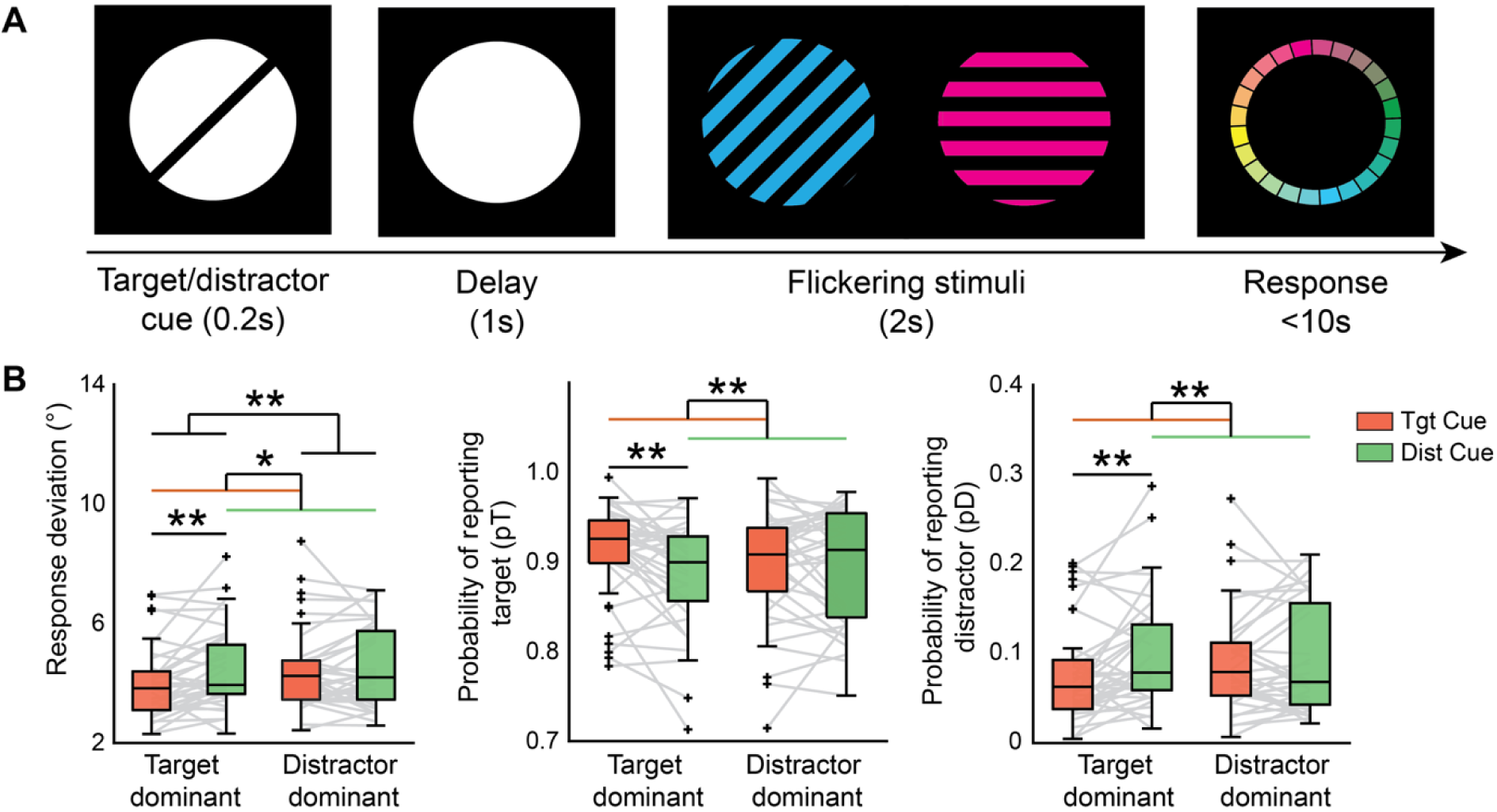
(A) Trial illustration. The target/distractor cue indicates the orientation of the corresponding grating. For half of the participants, target cues were in solid lines and the distractor cues were in dashed lines, while the rules were reversed for the other half participants. Gratings were presented to each eye using a stereoscope, with the target and distractor gratings flickering at 24 Hz and 20 Hz, respectively. During binocular rivalry, participants were instructed to prioritize the target grating while suppressing the distractor grating, aiming to reproduce the color of the target grating as precisely as possible. (B) Behavioral results. Response deviation represents how many degrees a response was off from the target color. From a probabilistic mixture model, we extracted the probability of reporting target color (pT), distractor color (pD) and random guess (pU) (Figure S1). Two-way ANOVA was performed on behavioral measurements. Results are shown for target cueing (TgtCue) and distractor cueing (DistCue) conditions, separately for when target was at the dominant eye versus when distractor was at the dominant eye. *, p < 0.05, **, p < 0.01, ***, p < 0.001. Probability of reporting target color, main effect of cueing: F(1, 140) = 9.939, p = 0.002, η² = 0.133; main effect of stimuli dominance: F(1, 140) = 3.452, p = 0.065; cueing x stimuli dominance, F(1,140) = 4.612, p = 0.033, η² = 0.098). Probability of reporting distractor color: main effect of cueing: F(1, 140) = 10.859, p = 0.001, η² = 0.154; main effect of stimuli dominance: F(1, 140) = 3.297, p = 0.072; cueing × stimuli dominance, F(1,140) = 4.370, p = 0.038, η² = 0.090).

## Results

### Absence of distractor-related cost in distractor-cueing trials

To assess how prior knowledge modulates perceptual competition, we first examined behavioral performance under target and distractor cueing. As expected, participants showed smaller color deviation in their responses when the target grating was presented to the dominant eye (target-in-dominant-eye trials; hereafter ‘target-dominant trials^1^’), compared to when distractor gratings were presented to the dominant eye (distractor-in-dominant-eye trials; hereafter ‘distractor-dominant trials’ for short, Figure 1B, F(1, 140) = 10.918, p = 0.001, η² = 0.199). Responses were more precise in target cueing condition than distractor cueing condition (F(1, 140) = 10.484, p = 0.002, η² = 0.157). Moreover, cueing effects differed as a function of stimuli dominance (F(1, 140) = 4.014, p = 0.047, η² = 0.084). In target-dominant trials, performance of the target cueing condition was better than distractor cueing condition (TgtCue vs. DistCue: p_bonferroni_ = 0.011). However, performance was comparable between the two cueing conditions in distractor-dominant trials (TgtCue vs. DistCue: p_bonferroni_ = 1.000).

To clarify which components of behavior contributed to this pattern, we applied a probabilistic mixture model on response deviations (Bays et al. 2009). The results showed that cueing modulated the likelihood of reporting the target and distractor in target-dominant trials, without affecting response precision or guessing rates (see Figure. S1). Specifically, participants were more likely to report the target color and less likely to report the distractor color in target cueing condition than distractor cueing condition when target was dominant (cueing effect on probability of reporting target: p_bonferroni_ = 0.004; cueing effect on probability of reporting distractor: p_bonferroni_ = 0.007). No such cueing effect was observed in distractor-dominant trials (p_bonferroni_ > 0.736).

### Feature cues boost sensory processing selectively at the dominant eye

The behavioral cueing effect in target-dominant trials could arise from two mechanisms. One possibility is that target cueing enhances the sensory representation or perceptual dominance of the target, either by stabilizing its dominance (Dieter and Tadin 2011) or reducing the competition from the distractor grating (Hancock and Timothy 2007), thereby improving performance. Under this account, distractor cueing would be expected to have little impact, as perceptual dominance already favors the target and the distractor is presented to the non-dominant eye. A key prediction of this mechanism is that distractor cueing should impair performance when the distractor occupies the dominant eye, unless such impairment is counteracted by inhibitory mechanisms.

An alternative account is that the cueing effect in target-dominant trials reflects a disruptive consequence of distractor cueing, whereby distractor cues bias initial selection toward the non-dominant distractor (Davidson et al. 2018; Ooi and He 1999), resulting in larger response deviation. This account predicts that prior cues are effective primarily when the cued stimulus is presented to the non-dominant eye. Accordingly, in distractor-dominant trials, where the target is non-dominant, target cueing should enhance perceptual dominance of the target and improve performance. However, this prediction was not supported by our behavioral results.

A third possibility is that the observed cueing effect reflects the combined influence of target facilitation and distractor capture. To distinguish between these alternatives and determine whether cueing selectively modulates target or distractor representations, we directly examined stimulus-specific sensory representations.

To determine how feature cues modulate sensory representations under visual competition, we examined orientation-selective neural responses to the two rivalrous gratings using multivariate decoding of posterior electrodes that were highly responsive to the flickering stimuli (Figure. 2A).

**Figure 2.**
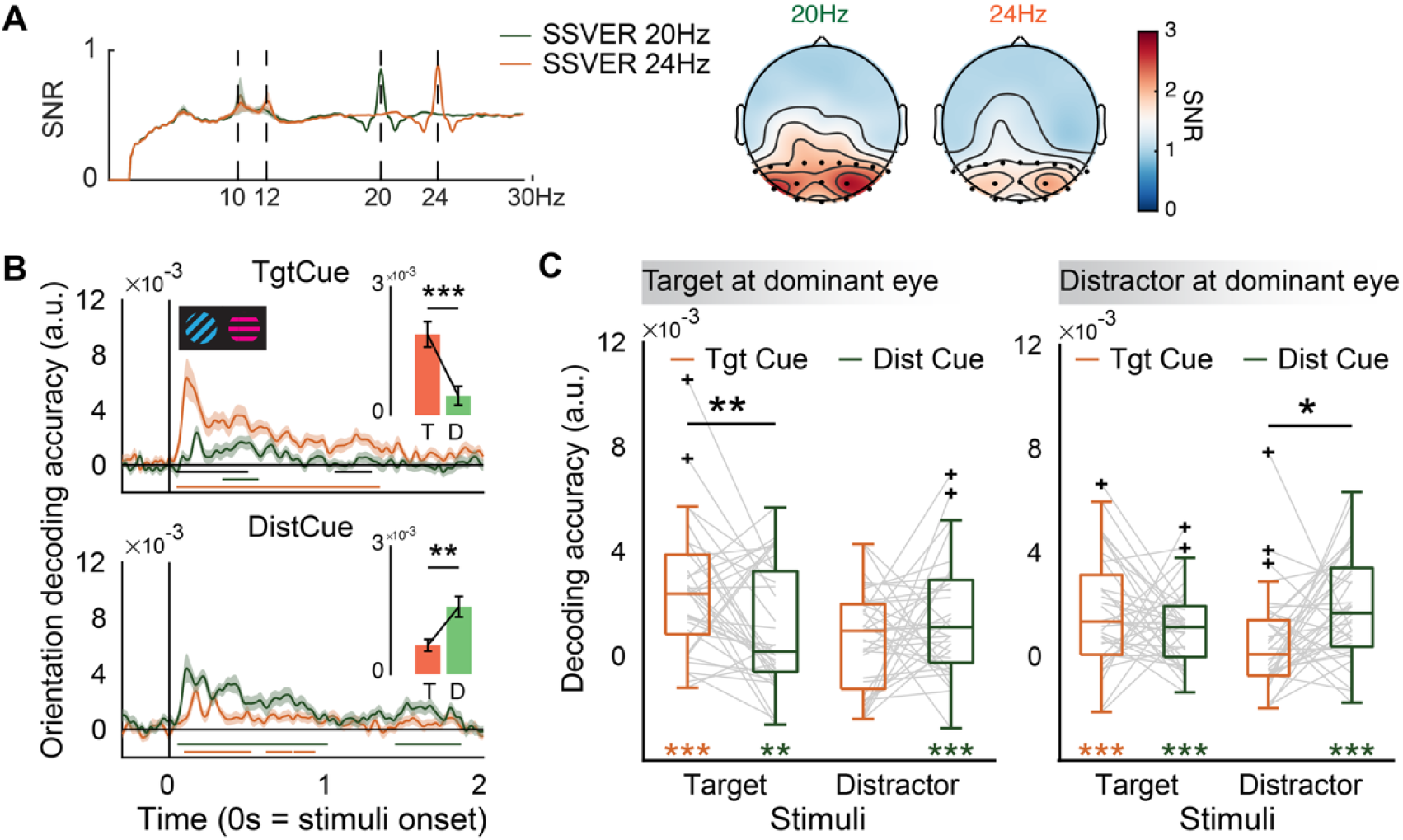
(A) Signal-to-noise ratio (SNR) of SSVER. The top panel shows normalized SNR using all channels. The topography highlights the most responsive channels to SSVER. (B) Mahalanobis-based distance decoding of target and distractor grating orientations in different cueing conditions. Colored lines above the x-axis represent time intervals with above-chance decoding accuracy, identified using cluster-based permutation testing. Black lines indicate the decoding accuracy difference between target and distractor gratings. Inserted panels show averaged decoding slopes calculated between 0 and 2 s during the rivalry phase. (C) Orientation decoding of target and distractor gratings based on stimulus dominance. Black asterisks mark significant differences between cueing conditions, while colored asterisks indicate significant decoding above chance level 0. For target dominant trials, target grating in TgtCue, t(35) = 5.916, p < 0.001, Cohen’s d = 0.986; distractor grating in TgtCue, t(35) = 1.852, p = 0.072, Cohen’s d = 0.309; target grating in DistCue, t(35) = 2.721, p = 0.010, Cohen’s d = 0.454; distractor grating in DistCue, t(35) = 3.887, p < 0.001, Cohen’s d = 0.648; For distractor-dominant trials, target grating in TgtCue, t(35) = 4.778, p < 0.001, Cohen’s d = 0.796; distractor grating in TgtCue, t(35) = 1.677, p = 0.103, Cohen’s d = 0.279; target grating in DistCue, t(35) = 4.611, p < 0.001, Cohen’s d = 0.769; distractor grating in DistCue, t(35) = 5.559, p < 0.001, Cohen’s d = 0.926.

Across trials, feature cues enhanced the orientation decoding accuracy of the cued stimulus, regardless of whether the cue referred to the target or the distractor (Figure. 2B, TgtCue: t(35) = 4.181, p < 0.001, Cohen’s d = 0.697; DistCue: t(35) = 3.053, p = 0.004, Cohen’s d = 0.509), indicating that cue-related biases in neural processing persist beyond the delay period. However, the cueing effect was further modulated by which eye the cued grating was presented to, with significant cueing effect primarily when it was shown to the dominant eye (Figure. 2C). When the target grating was presented to the dominant eye (Figure. 2C, left), orientation decoding of the target grating was significantly higher in target cueing compared to distractor cueing (t(35) = 2.815, p = 0.008, Cohen’s d = 0.469), whereas distractor grating did not differ between cueing conditions (t(35) = 1.605, p = 0.118). In contrast, when the distractor grating occupied the dominant eye, decoding of the distractor was enhanced by distractor cueing relative to target cueing (Figure 2C, right; t(35) = 2.714, p = 0.010, Cohen’s d = 0.452), with no corresponding modulation of target decoding (t(35) = 1.180, p = 0.246).

Together, these findings demonstrate that feature cues selectively boost sensory representations of the cued stimulus only when it is presented to the dominant eye. Importantly, this dominance-contingent pattern is inconsistent with the third account in which cueing effects arise from the combined but individually insignificant influences of target facilitation and distractor capture. Thus, the behavioral cueing effect observed in target-dominant trials reflects feature-based facilitation of the target representation.

### Dissociable cueing effects on alpha and theta activity during preparatory and rivalry phases

Although distractor cueing increased neural processing of the distractor in distractor-dominant trials, this enhancement did not translate into behavioral costs. This dissociation suggests that additional top-down control mechanisms counteracted distractor-related interference during perceptual competition. To identify the controlling mechanism, we examined neural rhythmic activities during the preparatory (pre-stimulus onset) and rivalry phases, focusing on theta (3-7Hz) and alpha (8-12 Hz) frequency bands.

During the preparatory phase, distractor cueing elicited stronger alpha power over right parietal regions compared to target cueing (Figure. 3A, t(35) = 2.152, p = 0.038, Cohen’s d = 0.359), whereas frontal theta power did not differ between cueing conditions during this period (Figure. 3B). If increased alpha power reflects direct suppression of the cued distractor feature, reduced neural representation of distractor orientation would be expected. However, orientation decoding revealed robust distractor representations during preparatory phase (Figure. 3C).

**Figure 3.**
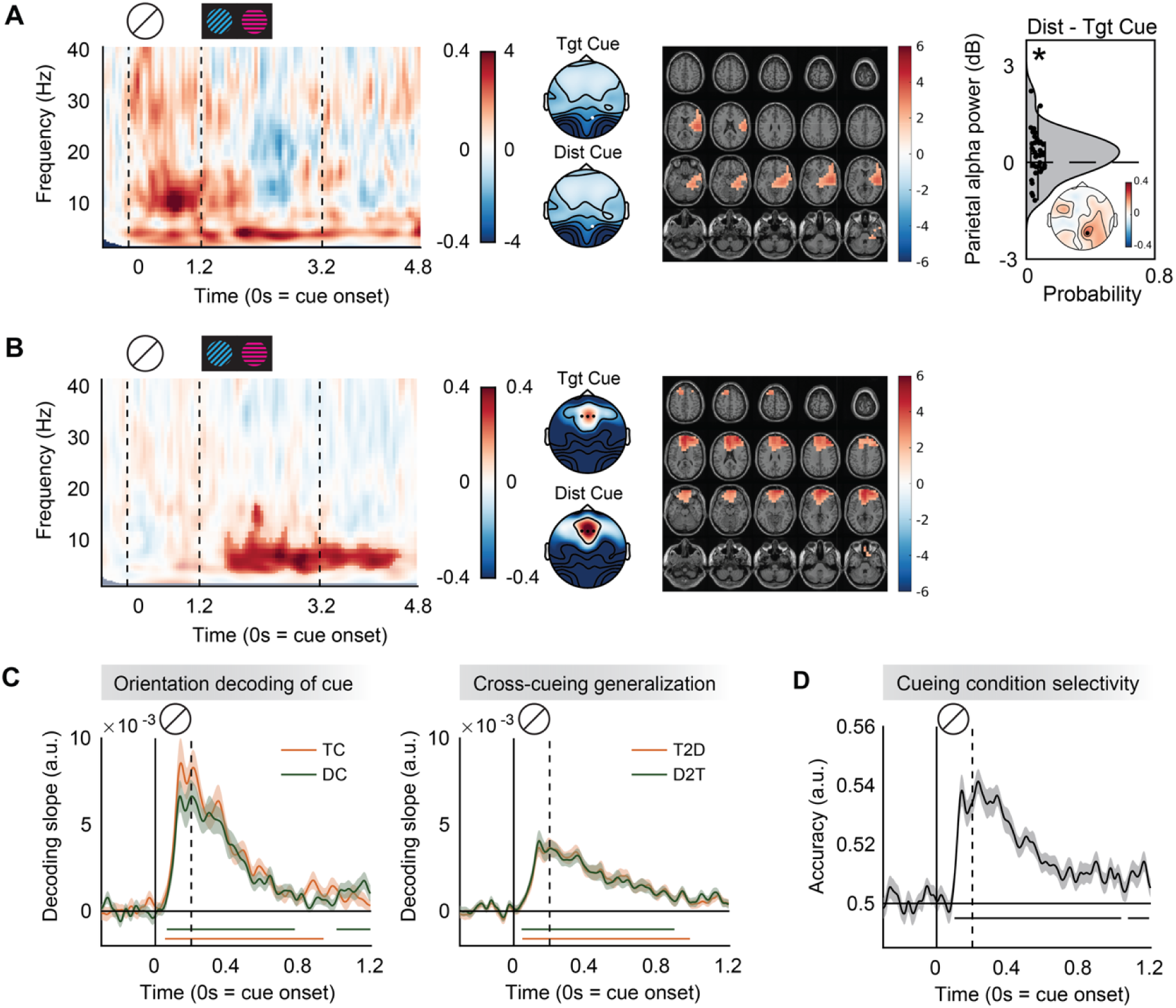
(A) Parietal alpha activity. The time-frequency map shows the power difference between target and distractor cueing conditions at electrode P2 (marked white). The right lateralized topographical distribution is consistent with previous findings showing the right posterior intraparietal sulcus in anticipatory alpha modulation (Capotosto et al. 2011). Source reconstruction revealed brain regions showed stronger alpha power (8-12 Hz) in distractor cueing condition compared to target cueing condition. Colorbar indicates t-value from cluster-based permutation test. The raincloud plot shows individual alpha power difference between cueing conditions. The inserted topography displays the averaged alpha power difference between cueing conditions during 0 to 1.2 s after cue onset. (B) Frontal theta activity. The time-frequency map shows the power difference between distractor and target cueing conditions, with the highlighted cluster indicating time-frequency points that passed cluster-based permutation testing. These results were derived from frontal channels highlighted in the topography. Source reconstruction revealed enhanced theta power in the distractor cueing condition during the rivalry phase. (C) Orientation decoding of cued orientations using posterior channels. Left, orientation decoding of each cueing condition separately; Right, robust cross-condition generalization (T2D: train target cueing data, test distractor cueing data; D2T: reverse) indicates that target and distractor templates utilize shared sensory-level representations. (D) Distinct task states. Multivariate classification of cueing conditions (TgtCue vs. DistCue) reveals distinct neural profiles throughout the delay, suggesting that the functional role of the template is maintained. The classifier was trained on data from all EEG.

Moreover, the orientation-based representations for target and distractor features were highly similar and could be generalized across cueing conditions (Figure. 3C right), indicating that preparatory alpha modulation did not attenuate feature-specific sensory representations. The generalizability of target and distractor representations is unlikely to reflect task indifference, as brain responses to target and distractor cues were reliably classifiable (Figure. 3D), indicating that the brain maintained distinct task states for the two cueing conditions. In addition, the persistence of strong distractor representations into the rivalry phase under distractor cueing further argues against early elimination of distractor features during preparation. Taken together, these findings argue against direct inhibition of distractor features during preparation, and instead suggest a feature-based gating mechanism that may shape upcoming sensory processing.

During the rivalry phase, distractor cueing was associated with significantly stronger low-frequency activity at the frontal region relative to target cueing (Figure. 3B, cluster-based permutation, 1.65 to 4.55 s, 3 to 16 Hz, p < 0.001; repeated ANOVA: main effect of dominance, F(1, 35) = 0.013, p = 0.911); main effect of cueing, F(1,35) = 19.515, p < 0.001, η² = 0.358, cueing x dominance, F(1, 35) = 0.344, p = 0.561), while parietal alpha power no longer differed between conditions. This temporal dissociation suggests a two-stage control process: preparatory alpha activity reflects proactive gating based on prior knowledge, whereas frontal theta activity during rivalry implements reactive control to suppress distractor interference as competition unfolds.

### Dissociable roles of alpha and theta in attentional control

To determine how the brain resolves distractor interference when sensory representations of the distractor are enhanced but behavior remains unaffected, we examined trial-by-trial relationships between neural oscillations, sensory gain, and performance using a general linear mixed model (see Table S1 for full model output and control analyses regarding theta during preparatory phase and alpha during rivalry phase).

Behavioral performance of each trial depended jointly on sensory evidence from both the target and the distractor (Figure. 4A left: target SSVER × distractor SSVER, F(1, 23147) = 8.463, p = 0.004). When target-related SSVERs were weak, stronger distractor SSVERs were associated with greater response deviations (F(1, 11595) = 4.962, p = 0.026), indicating increased interference (Figure. 4A right). This interference effect attenuated when target power was high (F(1, 11575) = 1.266, p = 0.261). Importantly, this target x distractor interaction effect was further modulated by cueing and stimulus dominance, particularly in the target cueing condition compared to the distractor cueing condition (target SSVER × distractor SSVER × cueing, F(1, 23147) = 9.251, p = 0.002), and especially when the target appeared in the dominant eye (target SSVER × distractor SSVER × cueing × stimuli dominance, F(1, 23147) = 7.751, p = 0.005). Thus, we separated target-dominant and distractor-dominant trials to examine the controlling mechanism.

**Figure 4.**
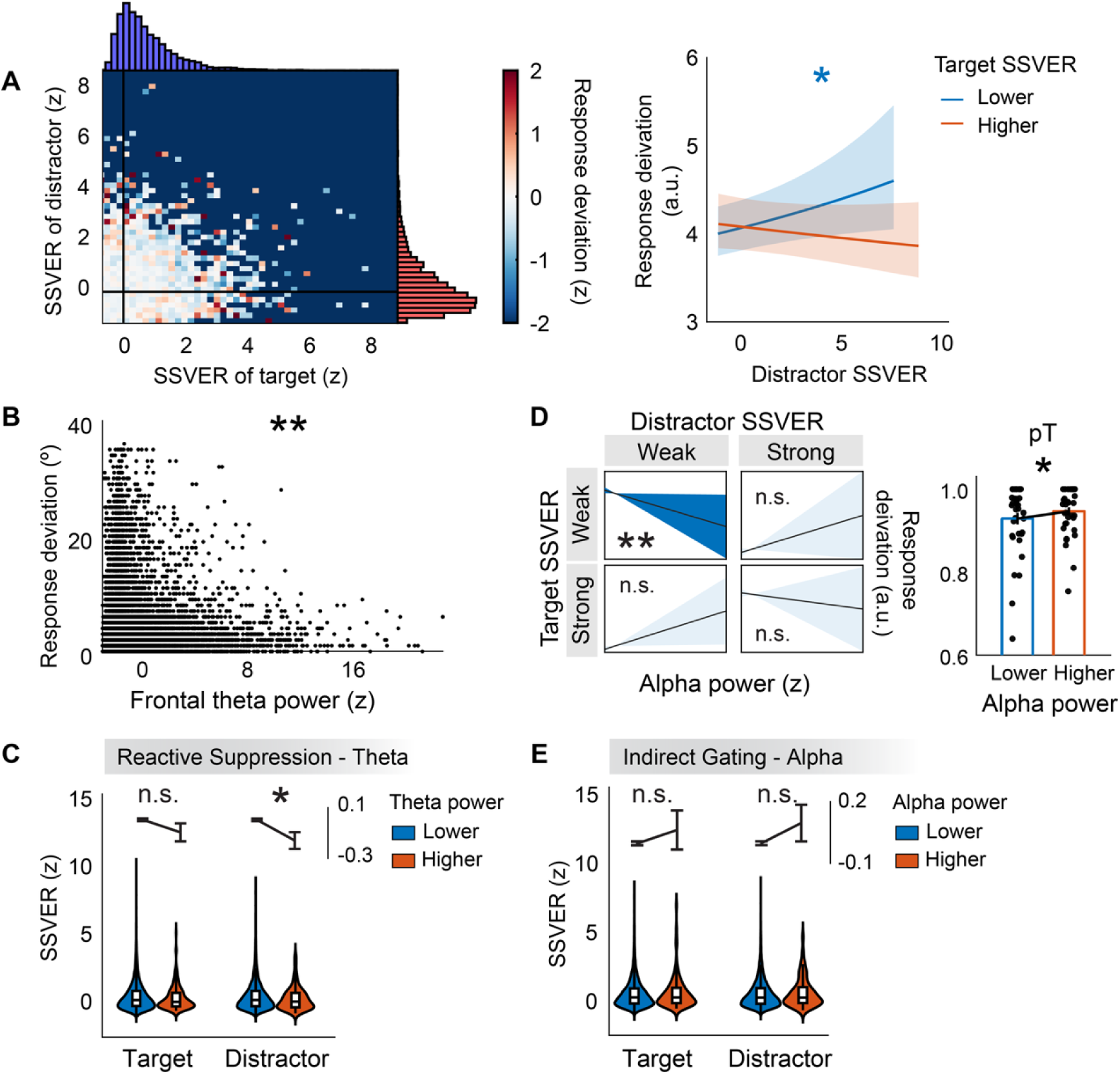
(A) Target SSVER and distractor SSVER jointly affect behavioral performance. Left, heatmap of response deviation (z-scored) as a function of target and distractor SSVERs (binned into 50 units) in target-dominant trials. The color of the scatter dots represents the average z-scores of behavioral response deviation within each bin. Dark regions indicate bins with insufficient trials. The top and right histograms show the probability density of target SSVER and distractor SSVER respectively (scale: 0 to 0.1). Right, linear regression coefficients for distractor SSVER in predicting deviation, median-split by target SSVER. Interference is significantly higher when target signals are weak (blue line). The intercept is not shown; the y-axis units are arbitrary reflecting relative values. (B-C) Frontal theta activity indexes reactive control associated with reduced distractor processing in target-dominant trials. (B) Increased frontal theta power (z-scored) correlates with reduced behavioral response deviation. (C) Stronger theta power is specifically associated with attenuated distractor SSVER, while target SSVER remains unaffected, suggesting a distractor-selective relationship between frontal theta and sensory processing. For visualization purpose, trials were median split based on frontal theta power of each participant to facilitate graphical interpretation of the effects. All statistical analyses were conducted on continuous single-trial data, and the categorical grouping was used solely for graphical presentation. (D-E) Parietal alpha power indexes indirect gating without altering sensory gain in distractor-dominant trials. (D) Under conditions of weak sensory gain (low SSVER), higher preparatory parietal alpha power is associated with smaller response deviations and an increased probability of target reporting (pT). (E) Preparatory alpha power does not directly modulate the SSVER of either target or distractor stimuli, supporting a role in higher-level gating rather than direct gain control.

Frontal theta activity during the rivalry phase selectively modulated the impact of sensory evidence on behavior in target-dominant trials (Figure. 4B: theta × target SSVER × distractor SSVER × stimuli dominance, F(1, 23147) = 4.128, p = 0.042). Higher theta power was associated with better performance, regardless of the SSVER of the target or distractor (main effect of theta power, F(1, 11578) = 7.112, p = 0.008). Follow-up analysis showed that increased theta power was accompanied by reduced distractor power, without significant changes in target power (Figure. 4C, distractor, t(11584) = 2.105, p = 0.035; target, t(11584) = 0.425, p = 0.671). These results indicate that frontal theta activity indexes reactive control associated with selective attenuation of distractor sensory processing during perceptual competition (Figure. 5 left).

**Figure 5.**
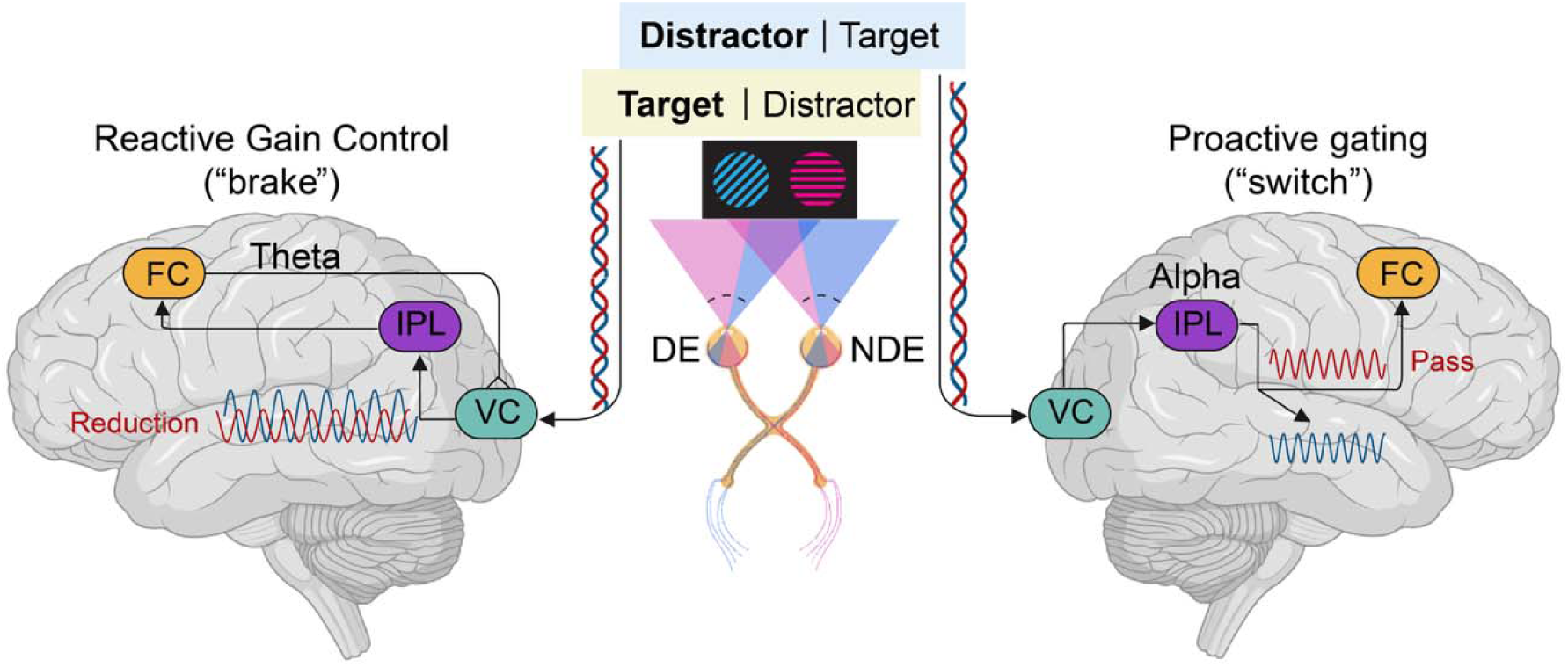
Reactive inhibition and proactive gating in resolving visual competition. Reactive inhibition via frontal theta (left panel): When the target is presented to the dominant eye, it receives a relative sensory advantage. Under this configuration, increased frontal theta is associated with reduced distractor sensory gain (red sinusoidal curve). This may help stabilize the target’s dominance thereby optimizing behavioral performance. Proactive attentional gating via parietal alpha (right panel): When the distractor occupies the dominant eye, high-uncertainty competition is resolved through proactive gating. Elevated preparatory alpha activity signals the instantiation of a negative attentional template based on prior distractor knowledge. This template serves to segregate and differentially route competing inputs without altering early sensory gain. This gating mechanism is most advantageous under conditions of high perceptual uncertainty due to weak sensory evidence, where it stabilizes weak target signals and facilitates the high-level processing. DE, dominant eye; NDE, nondominant eye; VC, visual cortex; IPL, inferior parietal lobe; FC, frontal cortex.

Preparatory parietal alpha activity modulated behavior in a qualitatively different manner (Figure. 4D: alpha × target SSVER × distractor SSVER × stimuli dominance, F(1, 23147) = 8.296, p = 0.004). Critically, in distractor-dominant trials, alpha power interacted with both target and distractor SSVERs (alpha × target SSVER × distractor SSVER, F(1, 11617) = 5.705, p = 0.017). To unpack this higher-order interaction, we categorized trials into combinations of relatively weak or strong target and distractor SSVERs. This follow-up analysis revealed that the alpha-related behavioral benefit was most pronounced under high perceptual uncertainty when both sensory signals were weak (Figure. 4D left, t(35) = 2.938, p = 0.006, Cohen’s d = 0.490).

Building on these results, we conducted an exploratory analysis to identify the specific cognitive component giving rise to the benefit. For each participant, trials in high perceptual uncertainty condition were further divided into higher-alpha and lower-alpha groups based on preparatory alpha power. Mixture models were then fitted to the two groups separately for each participant. We found that stronger preparatory alpha activity was associated with a higher probability of reporting the target under high perceptual uncertainty (Figure. 4D right, t(35) = 2.330, p = 0.026, Cohen’s d = 0.388), without affecting the precision or probability of reporting the distractor or random guesses (ps > 0.237, Figure. S2). This effect was not driven by any single participant (leave-one-participant-out validation, group differences = [0.018, 0.020], p = [0.0084, 0.0481]), and remained robust to participant resampling (bootstrap 95% CI for group difference = [0.003, 0.032]). This effect was most pronounced when both target and distractor SSVERs were weak, with no significant alpha modulation effect observed in the other three scenarios (Figure. S2, target weak-distractor strong, target strong-distractor weak, target strong-distractor strong).

Together, these findings suggest that increased preparatory alpha enhanced target identification and consequently facilitated behavioral performance, particularly under conditions of high perceptual uncertainty (Jensen 2024). In light of the behavioral influence of preparatory alpha activity, we further asked whether this alpha effect stemmed from direct modulation of sensory gain. To test this, we compared target and distractor SSVERs of trials with higher or lower preparatory alpha power. We failed to find alpha-related modulation of target SSVER (Figure. 4E, t(11623) = 0.458, p = 0.647) or distractor SSVER (t(11623) = 1.155, p = 0.248). Even when focusing on trials with weak target and distractor SSVER, alpha-related modulation effects on sensory gains were insignificant (target SSVER, t(3036) = 0.648, p = 0.517; distractor SSVER, t(3036) = 0.426, p = 0.670).

Together, these results revealed two dissociable control mechanisms operating at distinct stages of perception. Preparatory parietal alpha activity biased target selection through a gating process under weak sensory evidence, without altering sensory gain. In contrast, frontal theta activity during rivalry implemented reactive inhibition by suppressing distractor sensory gain. These dissociable processes provide a candidate account on how distractor representations can be enhanced at the sensory level without incurring behavioral costs.

We propose that elevated preparatory alpha reflects enhanced gating through the establishment of negative attentional templates informed by prior knowledge of the distractor (Carlisle 2023; de Vries et al. 2019; Jensen and Mazaheri 2010), allowing orientations mismatching the distractor to pass through (Figure. 5, right). During the feedforward processing of target and distractor, this template labels the two information streams and selectively passes the target inflow. Importantly, this separation and gating process does not alter the sensory gain of either the target or the distractor; instead, it routes them for further processing with different attentional weights. As a result, the target information is prioritized and stabilized, increasing its likelihood of being reported, particularly when sensory evidence is weak, thereby improving behavioral performance.

## Discussion

Here, we show that preparatory parietal alpha and rivalry-period frontal theta make dissociable contributions to attentional control at distinct stages of visual competition. Preparatory alpha was associated with gating based on prior distractor knowledge before stimulus onset, whereas frontal theta during rivalry was associated with reactive control and reduced distractor sensory processing. Together, these findings provide a candidate account of how prior information can support target-directed behavior despite enhanced sensory processing of anticipated distractors.

A central and initially counterintuitive finding of our study is that distractor cueing enhanced the sensory representation of distractors without incurring behavioral costs. This offers a possible explanation for an apparent paradox in the literature, where knowledge about distractors often captures attention rather than suppressing it (Cunningham and Egeth 2016; Geng et al. 2019; Noonan et al. 2016; Wen et al. 2018). Our data show that foreknowledge of a distractor equips the brain to dynamically counter this interference. During subsequent visual competition, frontal theta activity increased and was associated with better behavioral performance and selectively reduced distractor SSVER (Figure 5). This finding further demonstrates frontal theta not merely as a signal of conflict monitoring, but as an active "brake" limiting distractor-related processing (Cao et al. 2020; Cavanagh and Frank 2014; Messel et al. 2021). Given the broader involvement of frontal theta in cognitive and action control (Cooper et al., 2017; Panek et al., 2025), however, the present data cannot determine whether theta directly modulates distractor sensory processing or reflects higher-order cognitive or action control that subsequently influences sensory representations through top-down feedback.

Theta oscillations have frequently been linked to attentional sampling and rhythmic reorienting, particularly in tasks that require shifts of attention (Fiebelkorn and Kastner 2019; Galas et al. 2025; VanRullen 2016). Attentional shifting likely contributes to perceptual dynamics during binocular rivalry, where perceptual dominance alternates over time. However, several aspects of our data argue against a purely sampling-based account of theta activity in the present task. At the trial level, theta power was most strongly associated with improved behavioral performance when the target was presented to the dominant eye where attentional reorienting was expected to be least necessary. Moreover, stronger theta was observed when distractor representations were weaker, as indexed by decreased distractor-related SSVER responses. A shifting account would predict less attentional shifting when distractor is weak and stronger shifting when distractor is strong. These patterns are difficult to reconcile with a purely shifting-based account in which theta reflects the effort required to disengage from the cued or dominant distractor. Instead, they are more consistent with interpretations of theta as indexing reactive inhibitory control of distractors to establish stable perception, rather than rhythmic exploration or sampling of visual information.

In contrast to the reactive "brake" of theta, preparatory parietal alpha operated as a proactive "switch." Inconsistent with the "gain reduction" account of alpha-mediated inhibition, increased alpha power did not reduce the sensory gain of the distractor. Instead, our findings support the "gating-by-inhibition" hypothesis (Bonnefond and Jensen 2024; Peylo et al. 2021), where alpha oscillations functionally segregate competing inputs based on prior knowledge. This is supported by three key findings: (i) alpha power did not modulate the SSVERs of targets or distractors during the rivalry phase (Figure. 4E), (ii) in the distractor-cueing condition where alpha power was stronger, orientation selectivity for distractor gratings was enhanced rather than reduced (Figure. 2C), and (iii) stronger preparatory alpha was associated with increased probability of reporting target (Figure. 4D). We propose that this activity reflects the instantiation of a negative attentional filter tuned to the features of the expected distractor. By actively routing these matching signals away from higher-order processing centers, alpha protects the processing channel of the target, ensuring that limited cognitive resources are not squandered on irrelevant input. Notably, while attentional gating operates on every trial, it is most beneficial under high perceptual uncertainty where target and distractor signals are weak and difficult to distinguish. In these cases, template-based gating will compensate for the weak sensory input and facilitate the identification of target, thereby improving performance. In scenarios where one stimulus or both stimuli are strong, with prior information of target or distractor orientations, identifying target is unequivocal, additional gating conferred little benefit. This finding aligns with the perspective that alpha oscillations play a complementary role in indirect distractor inhibition, modulated by the load of goal-relevant information processing (Jensen 2024; Redding et al. 2026).

The sequential dynamics observed here could alternatively be considered in the context of a search-and-destroy account of distractor inhibition, in which an anticipated distractor is first represented to facilitate its detection and subsequently suppressed once encountered (Chelazzi et al., 2019; Moher & Egeth, 2012). Under this account, the distractor representation maintained during preparation could serve as a “search” template for identifying the distractor when the competing stimuli appear, followed by a reactive “destroy” stage during rivalry. Several aspects of our findings, however, do not readily conform to this interpretation. Preparatory distractor cueing increased parietal alpha while preserving a robust orientation-selective representation of the cued distractor, consistent with maintenance of a distractor template rather than its proactive suppression. Although this preserved representation could in principle constitute the search stage, the subsequent frontal theta effect does not show the pattern expected of a simple reactive destroy mechanism. Frontal theta was associated with better behavioral performance primarily when the target was presented to the dominant eye, rather than when the distractor was presented to the dominant eye, where a mechanism recruited to overcome a strongly competing distractor would be expected to have greater behavioral relevance. Moreover, the cued distractor representation remained decodable during rivalry, providing no evidence for wholesale elimination of its feature representation. Together, these findings favor a functional dissociation between preparatory gating and reactive control over a simple search-then-destroy sequence. Nevertheless, they do not exclude brief or transient episodes of distractor suppression during rivalry. Our trial-level SSVER measure was optimized to quantify sustained sensory gain rather than rapid within-trial dynamics, limiting our ability to resolve such transient changes. Search-and-destroy therefore remains a plausible alternative mechanism that cannot be definitively distinguished with the present design, but is not directly supported by the observed neural dynamics. Moreover, although binocular rivalry enabled stimulus-specific processing to be tracked during sustained visual competition, whether the dissociable contributions of preparatory alpha and rivalry-period frontal theta generalize to non-rivalrous forms of attentional competition remains an important question for future work.

Our findings also shed light on the nature of attentional templates and how they guide behavior. Both target and distractor cues in our study provided feature information that can be used to build attentional templates. Attentional templates can shape neural activity and affect attentional allocation (Carlisle 2023; Chelazzi et al. 2019; Woodman et al. 2013). An open question is whether target and distractor templates are represented as distinct templates or as a single content-based template with functional roles determined by task demands. The generalization of feature representations across target and distractor cueing conditions observed here supports the latter view. That is, target and distractor templates may share sensory content, while their functional consequences are determined by task context and control signals. The stronger parietal alpha activity observed during distractor cueing suggests that functional distinctions between templates, such as whether a feature should be selected or gated, may be encoded in non-sensory control signals rather than in the feature representation itself. This interpretation aligns with recent work on the dual-format model of attentional templates, which posits a dominant non-sensory format and a latent sensory-like format (Chen et al. 2024). Future studies comparing the dual-format representations of target and distractor templates will provide a more nuanced understanding of how attentional templates are maintained and utilized.

Beyond attentional control, our findings offer a mechanistic window into the neural signatures of conscious perception. Binocular rivalry has long served as a model system for studying the dynamics between conscious and unconscious states based on unchanging visual input (Seth and Bayne 2022; Tsuchiya et al. 2015). Pronounced endogenous neural oscillations in the theta-alpha range have been observed when experiencing mixed percepts (Cha and Blake 2019), or during perceptual switching (Davidson et al. 2018). Increased frontal theta activity before perceptual alternations is associated with resolving emerging conflicts, while increased alpha activity helps stabilizing perception through interocular inhibition (Drew et al. 2022). Our results extend this literature by showing that these rhythms play dissociable roles in shaping perceptual outcomes based on prior knowledge. According to Global Neuronal Workspace (GNW) theory, sensory inputs must be amplified and broadcast to a widespread network to reach awareness, which is known as ignition (Mashour et al. 2020; Seth and Bayne 2022). We found that alpha-mediated gating was most behaviorally beneficial when sensory evidence was weak (high perceptual uncertainty). In these near-threshold conditions, alpha-mediated routing improved the identification of the target, facilitating its ignition and subsequent entry into awareness. This suggests that parietal alpha plays a crucial role in the pre-conscious selection phase, acting as a spectral gatekeeper that determines which sensory streams are privileged for broadcast and which are silenced. Consistent with this view, stronger posterior alpha power has been reported in unseen correct trials than unseen incorrect trials (Trübutschek et al. 2017), and entraining posterior alpha activity selectively improves short-term memory performance for unseen stimuli that are visually unaware (Cheng et al. 2022). However, because we did not continuously track subjective perceptual experience during rivalry, eye-of-presentation indexed relative sensory advantage rather than moment-to-moment perceptual dominance, limiting direct inferences about subjective awareness. While future work is needed to directly test these ideas, from a broader theoretical perspective, our findings suggest that parietal alpha oscillations may contribute to ignition and broadcast, facilitating the routing of information necessary for conscious perception.

In conclusion, we demonstrate that resolving competing information involves two complementary mechanisms: reactive inhibition mediated by frontal theta oscillations and proactive attentional gating indexed by parietal alpha oscillations. This successive architecture explains how the brain maintains stable task-relevant perceptual processing despite the constant bombardment of distracting sensory information. By actively filtering the input (alpha) and reactively suppressing the noise (theta), the brain constructs a coherent visual reality from the chaos of rivalrous signals.

## Method

### Participants

A total of 38 participants were recruited to ensure a statistical power of 0.8 for detecting a medium effect size (d = 0.5) in a within-subject design, as determined by G*Power analysis. After preprocessing the electrophysiological data, two participants were excluded due to excessive trial rejection (38.8% of trials discarded), leaving a final sample of 36 participants (16 males; all right-handed) with an average age of 21 years (SD = 2.4, range: 18-26 years). Eight participants exhibited left-eye dominance. All participants had normal or corrected-to-normal vision, normal color vision, and no reported history of psychiatric or neurological disorders. Participants completed a training session on the first day to familiarize themselves with the experimental tasks and confirm the occurrence of the binocular rivalry effect. On the second day, scalp electrophysiological recording was conducted. Written informed consent was obtained from all participants before the experiment. The study was approved by the Institutional Review Board of the School of Psychological and Cognitive Sciences at Peking University (Protocol No. 2020-02-02).

### Task

The experiment was conducted in a dimly lit, electromagnetically and acoustically shielded room. Visual stimuli were presented on a gray background using a Display++ monitor (refresh rate: 120 Hz, resolution: 1920 × 1080) and viewed through a mirror stereoscope to ensure independent stimulus presentation to the two eyes. Ocular dominance was assessed for each participant before the experiment using the hole-in-the-card test, and stimulus assignment was then determined according to the participant’s dominant eye. Each trial began with a cue displayed for 0.2 seconds (length = 4°, width = 0.2°) to indicate the orientation of the target or the distractor. Target cueing and distractor cueing conditions were consistent within a block and were indicated using solid lines and dashed lines. Participants were informed of the cueing condition before each block. The line type (solid or dashed) and its association with the target or distractor cueing was counterbalanced across participants. Participants were informed of the association prior to the experiment. After a 1 s delay, two gratings of different colors and orientations were presented separately to each eye for 2 s (diameter: 10°, spatial frequency: 0.04 cycles/pixel, contrast = 1). We selected the stimulus duration based on a previous study showing a mean dominance duration for about 2 seconds (Zhang et al. 2011). The short duration limited natural perceptual switching. Thus, performance was likely to be biased toward the distractor if it captured the initial dominance. One grating’s orientation matched the cued orientation. In the target cueing condition, the orientation-matched grating served as the target, whereas in the distractor cueing condition, the non-matching grating became the target. Participants were required to recall the target grating’s color and report it by selecting the corresponding hue on a color wheel within 10 seconds, with an emphasis on precision. Participants completed 720 trials across 16 blocks, with 45 trials per block. The cueing condition was manipulated between blocks to avoid task switch cost.

A no-cue condition was not included in the present experiment. In the context of binocular rivalry, the absence of a cue would require participants to attend both stimuli simultaneously in order to perform the task, imposing an attentional load of two. Such a condition would therefore introduce a qualitatively different attentional state that is not directly comparable to target-cueing or distractor-cueing conditions, in which attentional priorities are explicitly specified and therefore an attention load of one. Moreover, in a no-cue condition, prolonged mixed percepts are more likely to occur, reflecting unresolved visual competition rather than active competition resolution. This would confound the interpretation of neural signatures related to attentional gating and suppression. Although the absence of a no-cue baseline precludes direct behavioral comparison against the baseline, the within-subject contrasts between cueing and dominance are sufficient to isolate the effects of attentional guidance on perceptual competition.

The orientations of the gratings were selected from six orientation bins, with the angular difference between the target and distractor gratings always being at least 15° to enhance visual discrimination. Grating colors were randomly selected for each trial from a set of 72 colors in the Lab color space (L = 70; a*, b* plane origin at [0, 0]) to ensure uniform perceived brightness. The angular difference between the colors of any two gratings was a minimum of 60°.

To enhance competition between the two eyes, the phase of the gratings was reversed at different intervals: every 5 frames (24 Hz) for the target grating and every 6 frames (20 Hz) for the distractor grating. This manipulation compelled participants to selectively attend to the target grating while inhibiting the distractor. Participants were instructed to fixate on the center of the screen and minimize blinking, as eye movements and blinks can disrupt binocular rivalry.

Behavioral effects were assessed using linear mixed-effects models with cueing condition, stimulus dominance, and their interaction as fixed effects, and a random intercept for each participant. Partial eta squared (η²) was calculated from corresponding repeated-measures ANOVAs with the same experimental factors to provide complementary effect-size estimates.

### EEG recording and preprocessing

The EEG was recorded using Ag/AgCl electrodes placed on a 64-channel EasyCap according to the international 10-10 system (Brain Products). An external electrode placed below the right eye was used to measure eye movements. Electrode impedance was kept below 10 kΩ. The data were collected at a sampling rate of 1000 Hz. Offline preprocessing was performed using the EEGLAB toolbox (Delorme and Makeig 2004). Raw data were downsampled to 500 Hz and re-referenced using the average of bilateral mastoid electrodes. A bandpass filter of 1 to 40 Hz was applied. Bad channels were spherically interpolated. The continuous data were epoched into -1 s to 4 s time locked to cue onset. Typical artifacts such as blinks, eye movements, heartbeat noise, muscle noise, and line noise were identified and corrected using independent component analysis. There were on average 4.4 (SD = 1.6) components removed. After preprocessing, trials containing voltage values exceeding a threshold of 100 μV at any time point for any electrode were excluded. After preprocessing, 94.1% of trials on average were retained across participants (SD = 4.09%; range = 80.1-98.2%). For subsequent EEG analyses, trials with extreme behavioral response deviations were additionally excluded using a participant-specific threshold defined as the smaller of 45° and the 95th percentile of the participant’s absolute target-response deviation. This behavioral criterion excluded an additional 4.75% of the remaining trials on average (SD = 0.32%; range = 3.43-5.07%).

### SSVER extraction

We use Rhythmic Entrainment Source Separation (RESS) to isolate SSVERs induced by the flicking gratings (Cohen, Michael X. and Gulbinaite 2017). This method combines and extends existing multivariate source separation methods to optimize the extraction of steady-state responses from EEG signals. The full width at half maximum (FWHM) of the peak frequency was 0.5 Hz, and the distance of neighboring frequencies away from peak frequency was 1 Hz, the FWHM of the neighboring frequencies was 1 Hz. The time interval was selected to be 1.3 s to 3.1 s after cue onset to avoid transient stimulus-evoked effect. Note that while there were strong sub-harmonics in alpha band at 10Hz and 12Hz, our analysis on alpha activity was at the preparatory phase before the onset of flickering stimuli. These sub-harmonics could not have confounded our findings regarding proactive alpha-gating.

### Time frequency analysis

Time-frequency analysis was conducted on the entire epoch (-1 to 4s time locked to cue onset) using a Morlet wavelet decomposition in the FieldTrip toolbox (Oostenveld et al. 2011). The decomposition was implemented by specifying a range of frequencies of interest from 1 to 40 Hz on a combined linear scale (1-40 Hz) with a logarithmic scale to achieve finer frequency resolution at lower frequencies. The wavelet width was linearly scaled from 2 to 10 cycles across frequencies. Time points of interest were selected every 50 ms across the trial window. Normalization was carried out using decibel transformation, with the baseline interval set from - 0.4 to -0.1 s relative to the pre-cue period. To ensure that our results reflected fluctuations in endogenous oscillatory amplitude rather than ERP differences between conditions, all time-frequency analyses were performed on non-phase-locked data. Specifically, ERP components were removed by subtracting the condition-specific ERP from the single-trial EEG data before performing the time-frequency decompositions on the residual data (Cohen, Mike X. 2014). Cueing-related differences in time-frequency power were assessed using cluster-based permutation testing across the full 1-40 Hz range, without restricting the analysis to predefined frequency bands.

### Source reconstruction

To investigate the cortical sources of theta activity during the rivalry and alpha activity during the preparatory phase, we performed source reconstruction using the Dynamic Imaging of Coherent Sources (DICS) beamforming method. After computing the forward model using standard MRI, the cross-spectral density matrix was computed from the averaged theta (3-7Hz) and alpha (8-12 Hz) frequency bands, derived via a multitaper fast Fourier transform. To account for the increased noise towards the center of the head, the neural activity index (NAI) was calculated for each voxel in the brain by dividing the source estimates by the corresponding local noise estimates. Power values of the theta and alpha frequency bands were extracted from 0.3 to 1.2 s after stimuli onset and 0.6 to 1.1 s after cue onset respectively. The statistical significance of clusters showing differences between target cueing and distractor cueing conditions was assessed by Monte Carlo permutation test (N = 10000) with two-tailed alpha of 0.05.

### Orientation decoding

We performed orientation decoding using data from 17 posterior channels. Data was partitioned into seven training folds and one test fold for cross-validation. To ensure an unbiased classifier, we subsampled equalized number of trials for each orientation bin in the training set and then took the trial-average of each orientation bin. After obtaining the covariance matrix across orientation bins of the training set, we calculated the Mahalanobis distances between the test trials and the training sets (Wolff et al. 2017). A smaller distance toward one specific orientation bin indicated a larger pattern similarity. Decoding accuracy was measured as the cosine weighted distances of pattern-difference between trials of increasingly dissimilar orientations. The cross-validation procedure was iterated 100 times, with random assignment of trials to the training and test sets in each iteration. The averaged decoding accuracy was taken for the statistical analysis. We followed the same procedure for cross-condition decoding except that the model was trained from one cueing condition (e.g., target cueing) and tested on the other condition (e.g., distractor cueing).

Finally, to assess the statistical significance of the decoding accuracy, we conducted cluster-based permutation testing against the chance level (Maris and Oostenveld 2007), using an alpha threshold of 0.05. Cluster statistics were calculated using the sum of the values within each cluster, and the permutation test was run for 10,000 iterations to generate the null distribution for comparison. The cluster-based nonparametric alpha was set to 0.05.

### General linear mixed model on trialwise behavioral and neural responses

We conducted a general linear mixed model to analyze trialwise behavioral and neural responses. The dependent variable (trial-wise response error) is predicted by the fixed effects (cueing condition, stimuli dominance, reactive frontal theta power, preparatory parietal alpha power, target and distractor SSVER strength, and their interactions), and the random effects between. The model formula was: beha ∼ alpha*theta*target power*distractor power*cueing condition*stimuli dominance + (1|subjects). Behavioral color deviation for each trial was the outcome variable. Preparatory parietal alpha power was extracted from P2 based on the cueing-effect topography, reactive frontal theta power was extracted from Fz, SSVERs for target and distractor at tagging frequencies were extracted from PO3 and PO4. Neural measurements were normalized within each participant before entering the model. The response distribution was specified as “normal,” and the link function as “identity.”

Model convergence was verified using MATLAB’s penalized-likelihood optimization diagnostics. The relative change in the linear predictor at convergence was well below the specified tolerance, and the estimated participant-level random-effect covariance parameter was finite and away from the boundary. In addition to the random-intercept model, we fitted two more complex models that included the key within-subject experimental factors, cueing condition and eye-of-presentation, in the random effect. While the log-likelihood increased modestly with additional random-effects parameters, this improvement was insufficient to offset the added model complexity. Both AIC and BIC favored the random-intercept model.

After identifying condition-specific modulations of alpha and theta power, we examined whether these oscillatory signals influenced the sensory gain of target and distractor. To this end, we performed single-trial GLMM where stimulus SSVERs were predicted by trial-wise alpha or theta power: stimulus SSVER ∼ alpha (or theta) + (1 | subject). Oscillatory power was entered as a continuous predictor and subjects were modeled with random intercepts to account for between-participant variability. For visualization purposes only (Figure 4), trials were median-split within each participant into high-and low-alpha (or theta) subgroups. This binning was used solely to facilitate graphical interpretation of the effects and was not applied in any statistical analyses.

To assess the robustness of the alpha-related behavioral effect, we performed leave-one-participant-out and bootstrap analyses. Within each participant, trials with relatively weak target and distractor SSVERs were divided into low- and high-alpha groups based on the median preparatory alpha power. Mixture model was fitted to each alpha group separately to extract response precision, probability of reporting target color, distractor color and random guess. For the leave-one-participant-out analysis, we repeated the paired-samples t-test 36 times, excluding one participant in each iteration. We recorded the range of effect estimates and associated p-values to determine whether the effect depended on any single participant. For the bootstrap analysis, we resampled participants with replacement while preserving their paired high- and low-alpha estimates. We calculated the mean high-minus-low difference for each of 1,000 bootstrap samples and derived the 95% confidence interval from the 2.5th and 97.5th percentiles of the resulting distribution.

## Data availability

Data and script can be found at OSF at https://osf.io/3dnvy/

## Acknowledgments

We thank Chenyang (Leo) Lin for constructive discussions and Yangming Zhang for assistance with data collection. This study was supported by grants from the National Natural Science Foundation of China (32271104) and STI2030-Major Projects (2021ZD0200204).

## Author contributions

WW and SL designed the study. WW and YW collected data. WW analyzed the data. WW wrote the manuscript. WW, RR, and SL revised the manuscript.

## Footnotes

1 Because participants were not required to report the dominant percept, we cannot directly determine which stimulus held perceptual dominance at a given moment. Instead, trials were categorized based on whether the target or distractor grating was presented to the participant’s dominant eye. While stimuli presented to the dominant eye typically gain initial perceptual dominance, this is not guaranteed on every trial and may not persist throughout the entire trial. Unless otherwise noted, the terms “target-dominant” and “distractor-dominant” refer specifically to stimulus presentation to the dominant eye, rather than the perceptual experience.

