## Supplementary figures and table for "Alpha and theta dynamics successively resolve competition between rival visual stimuli"

### Figure. S1

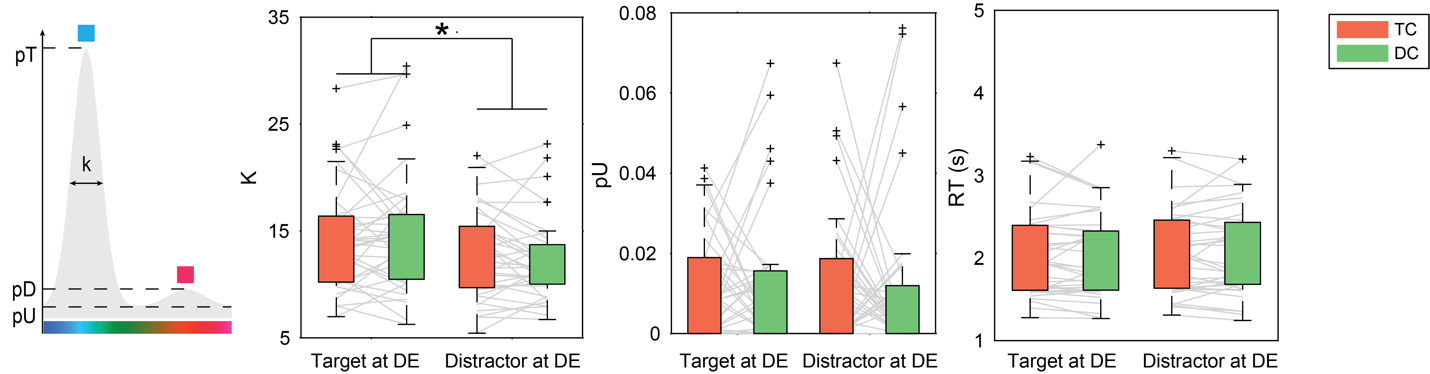

Fig. S1. Mixture model and behavioral responses. In the probability mixture model, k represents the response precision, pT and pD represent the probability of reporting target and distractor feature respectively, pU represents the probability of a random guess (Bays et al., 2009). Color precision was significantly better when the target grating was presented to the dominant eye compared to when distractor was in the dominant eye (F(1, 140) = 6.705, p = 0.011, η²ₚ = 0.340), and this effect was independent of cueing conditions (cueing: F(1, 140) = 0.167, p = 0.683; cueing × stimuli dominance: F(1, 140) = 0.865, p = 0.354). Participants showed similar guess rates across conditions (stimuli dominance: F(1,140) = 0.140, p = 0.709; cueing: F(1,140) = 0.060, p = 0.806; cueing × stimuli dominance: F(1,140) = 0.202, p = 0.654). Reaction time did not differ significantly across conditions (stimuli dominance: F(1,140) = 0.385, p = 0.536; cueing: F(1,140) = 0.075, p = 0.784; cueing × stimuli dominance: F(1,140) = 0.355, p = 0.552).

### Figure. S2

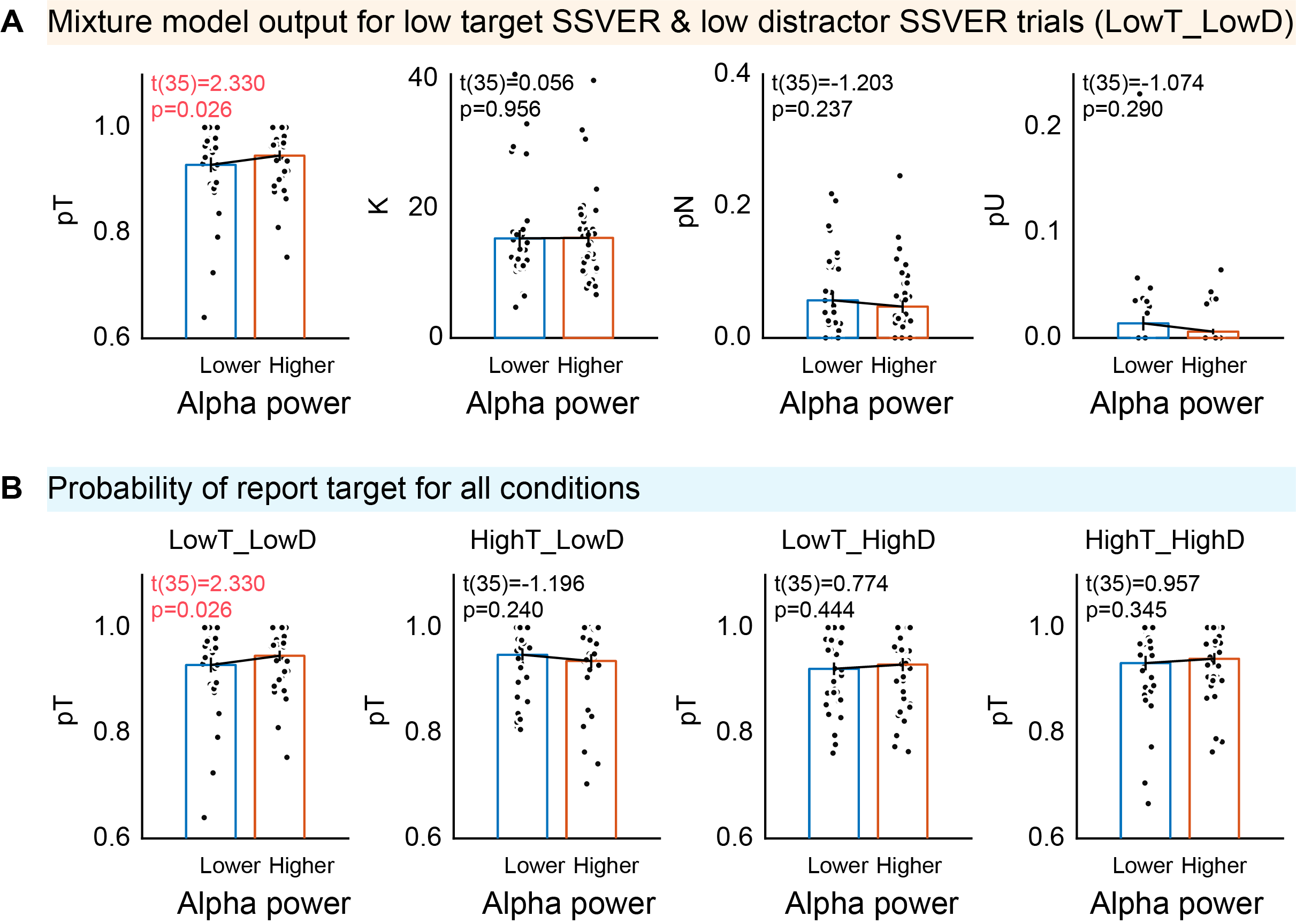

Figure S2. Behavioral probability mixture model output for all trial combinations. (A) For trials with lower target power and lower distractor power, stronger preparatory alpha led to increased probability of reporting target, but no significant modulation effect on probability of reporting distractor or random guess, nor response precision. (B) Alpha modulated probability changes in reporting target were only observed in trials with lower target and distractor input.

### Figure. S3

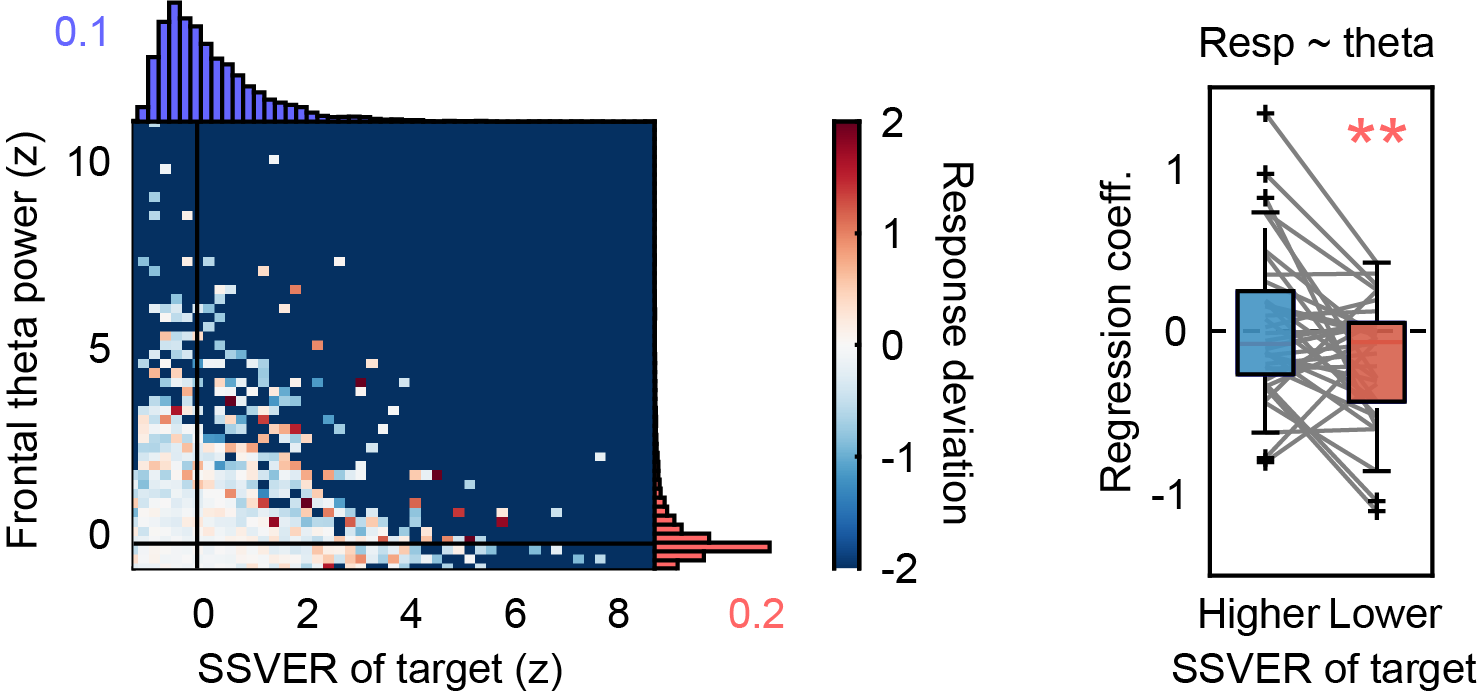

Fig. S3. Behavioral performance modulated by theta power and target SSVER. There was a significant interaction effect on Theta × Target × Cue × Dominance (F(1, 23147) = 4.750, p = 0.029). Parsing out the four-way interaction, we found that in the target cueing condition when target appeared at the dominant eye, whereas stronger frontal theta power would enhance performance leading to reduced response deviation when target SSVER was low (t(35) = 3.032, p = 0.005, Cohen’s d = 0.505), but not when target SSVER was high (t(35) = 0.443, p = 0.661).

### Table. S1

| **Term** | **FStat** | **DF1** | **DF2** | **pValue** |
| --- | --- | --- | --- | --- |
| (Intercept) | 257.7 | 1 | 23147 | 1.12E-57 |
| Theta | 3.722 | 1 | 23147 | 0.053708 |
| Alpha | 0.224 | 1 | 23147 | 0.63568 |
| Target | 0.91 | 1 | 23147 | 0.34011 |
| Distractor | 1.284 | 1 | 23147 | 0.25726 |
| Cue | 0.771 | 1 | 23147 | 0.38004 |
| Dom | 22.408 | 1 | 23147 | 2.22E-06 |
| Theta:Alpha | 0.027 | 1 | 23147 | 0.87023 |
| Theta:Target | 4.521 | 1 | 23147 | 0.033487* |
| Alpha:Target | 0.753 | 1 | 23147 | 0.38556 |
| Theta:Distractor | 2.036 | 1 | 23147 | 0.1536 |
| Alpha:Distractor | 1.703 | 1 | 23147 | 0.19189 |
| Target:Distractor | 8.463 | 1 | 23147 | 0.003628 |
| Theta:Cue | 0.001 | 1 | 23147 | 0.97703 |
| Alpha:Cue | 0.207 | 1 | 23147 | 0.64917 |
| Target:Cue | 0.034 | 1 | 23147 | 0.85451 |
| Distractor:Cue | 0.556 | 1 | 23147 | 0.45575 |
| Theta:Dom | 0.432 | 1 | 23147 | 0.51122 |
| Alpha:Dom | 0.999 | 1 | 23147 | 0.3176 |
| Target:Dom | 3.38 | 1 | 23147 | 0.066012 |
| Distractor:Dom | 0.301 | 1 | 23147 | 0.58354 |
| Cue:Dom | 1.751 | 1 | 23147 | 0.18579 |
| Theta:Alpha:Target | 1.091 | 1 | 23147 | 0.29617 |
| Theta:Alpha:Distractor | 0.484 | 1 | 23147 | 0.48666 |
| Theta:Target:Distractor | 1.432 | 1 | 23147 | 0.23139 |
| Alpha:Target:Distractor | 3.435 | 1 | 23147 | 0.063854 |
| Theta:Alpha:Cue | 0.139 | 1 | 23147 | 0.70893 |
| Theta:Target:Cue | 2.154 | 1 | 23147 | 0.14218 |
| Alpha:Target:Cue | 1.414 | 1 | 23147 | 0.23437 |
| Theta:Distractor:Cue | 1.929 | 1 | 23147 | 0.16494 |
| Alpha:Distractor:Cue | 0.366 | 1 | 23147 | 0.54533 |
| Target:Distractor:Cue | 9.251 | 1 | 23147 | 0.0023563* |
| Theta:Alpha:Dom | 0.047 | 1 | 23147 | 0.82842 |
| Theta:Target:Dom | 5.223 | 1 | 23147 | 0.022296* |
| Alpha:Target:Dom | 0.104 | 1 | 23147 | 0.7471 |
| Theta:Distractor:Dom | 0.462 | 1 | 23147 | 0.4969 |
| Alpha:Distractor:Dom | 0.461 | 1 | 23147 | 0.49722 |
| Target:Distractor:Dom | 3.471 | 1 | 23147 | 0.062465 |
| Theta:Cue:Dom | 1.544 | 1 | 23147 | 0.21405 |
| Alpha:Cue:Dom | 0.257 | 1 | 23147 | 0.61232 |
| Target:Cue:Dom | 0.655 | 1 | 23147 | 0.41833 |
| Distractor:Cue:Dom | 0.252 | 1 | 23147 | 0.61546 |
| Theta:Alpha:Target:Distractor | 1.065 | 1 | 23147 | 0.30206 |
| Theta:Alpha:Target:Cue | 0.839 | 1 | 23147 | 0.35963 |
| Theta:Alpha:Distractor:Cue | 0.789 | 1 | 23147 | 0.37442 |
| Theta:Target:Distractor:Cue | 2.308 | 1 | 23147 | 0.12874 |
| Alpha:Target:Distractor:Cue | 2.255 | 1 | 23147 | 0.13316 |
| Theta:Alpha:Target:Dom | 0.005 | 1 | 23147 | 0.94521 |
| Theta:Alpha:Distractor:Dom | 0.702 | 1 | 23147 | 0.40228 |
| Theta:Target:Distractor:Dom | 4.128 | 1 | 23147 | 0.042197* |
| Alpha:Target:Distractor:Dom | 8.296 | 1 | 23147 | 0.0039774* |
| Theta:Alpha:Cue:Dom | 0.332 | 1 | 23147 | 0.56478 |
| Theta:Target:Cue:Dom | 4.75 | 1 | 23147 | 0.029311* |
| Alpha:Target:Cue:Dom | 0.069 | 1 | 23147 | 0.79262 |
| Theta:Distractor:Cue:Dom | 1.191 | 1 | 23147 | 0.27506 |
| Alpha:Distractor:Cue:Dom | 0.779 | 1 | 23147 | 0.37743 |
| Target:Distractor:Cue:Dom | 7.751 | 1 | 23147 | 0.0053718* |
| Theta:Alpha:Target:Distractor:Cue | 1.506 | 1 | 23147 | 0.21985 |
| Theta:Alpha:Target:Distractor:Dom | 0.27 | 1 | 23147 | 0.60357 |
| Theta:Alpha:Target:Cue:Dom | 0.119 | 1 | 23147 | 0.73026 |
| Theta:Alpha:Distractor:Cue:Dom | 1.578 | 1 | 23147 | 0.20904 |
| Theta:Target:Distractor:Cue:Dom | 2.883 | 1 | 23147 | 0.089539 |
| Alpha:Target:Distractor:Cue:Dom | 2.186 | 1 | 23147 | 0.13933 |
| Theta:Alpha:Target:Distractor:Cue:Dom | 0.037 | 1 | 23147 | 0.84649 |

Full results of the general linear mixed model. Alpha, alpha power (10Hz) obtained from P2 during the preparatory phase; Theta, theta power (6Hz) obtained from Fz during the rivalry phase; Target, 24Hz SSVER of PO3 and PO4; Distractor, 20Hz SSVER of PO3 and PO4. Cue, cueing conditions; Dom, stimuli dominance. Channel of interest was selected based on topographical distribution. Significant interaction effect on Theta × Target × Cue × Dom was illustrated in Fig. S3. The observed effects were robust even after controlling the influence of parietal alpha power during the rivalry phase and frontal theta power during the preparatory phase. When these factors were included as random effects in the general linear mixed model, the critical four-way interactions remained significant. Theta x Target x Distractor x Dom, F(1, 23147) = 4.126, p = 0.042; Alpha x Target x Distractor x Dom, F(1, 23147) = 8.307, p = 0.004; Targe x Distractor x Cue x Dom, F(1, 23147) = 7.749, p = 0.005. Theta x Target x Cue x Dom, F(1, 23147) = 4.751, p = 0.029.
